# Serotonergic Effects on Interoception

**DOI:** 10.1101/2020.08.28.262550

**Authors:** James J A Livermore, Clare L Holmes, Gyorgy Moga, Kristian Adamatzky, Hugo D Critchley, Sarah N Garfinkel, Daniel Campbell-Meiklejohn

## Abstract

Interoception is the signalling, perception, and interpretation of internal physiological states. Much of the psychopharmacology of interoception is still undiscovered. However, psychiatric disorders associated with changes of interoception, including depressive, anxiety, and eating disorders are often treated with selective serotonin reuptake inhibitors (SSRIs). The causal effect of acute changes of serotonin transmission on interoceptive cognition was tested by a within-participant, crossover, placebo-controlled study. Forty-seven healthy human volunteers (31 female, 16 male) were tested both on and off a 20mg oral dose of the commonly prescribed SSRI citalopram. For each randomly ordered session, participants made judgments on the synchrony of their heartbeat to auditory tones and expressed confidence in each of these judgments. Citalopram enhanced insight into the likelihood that one’s interoceptive judgment had been correct, driven primarily by enhanced confidence for correct responses. This effect was independent of measured cardiac and subjective effects of the drug. This novel result is evidence that acute serotonin changes can alter metacognitive insight into the reliability of inferences based on interoceptive information, which is a foundation for considering effects of serotonin on cognition and emotion in terms of effective top-down regulation of interoceptive influence on mental states.

## Introduction

Interoception is the primary driver of allostasis, through which our health and vitality is maintained by dynamic, and often predictive, adaptations in our physiology, cognition, and behavior. By signaling, perceiving, and interpreting our internal physiological states, interoception changes how we feel, what we choose, how we react, what we learn, our physiological sense of ‘self’ and our beliefs about our situation [1–3]. Little is known about the psychopharmacology of interoceptive cognition. However, interoceptive changes are now known to feature as a transdiagnostic feature across psychiatric conditions, including depressive, anxiety, and eating disorders [4–7], a majority of which are treated with selective serotonin reuptake inhibitors (SSRIs). To our knowledge, a causative relationship between acute serotonin changes and interoception has never been reported outside of the sensory domain of pain, making the present study necessary to bridge our understanding of interoception to neurocognitive models of antidepressant action in psychiatric disorders (e.g. [8]). In this study, we tested the relationship between acute changes in serotonin signalling and cardiac interoception.

Selective serotonin reuptake inhibitors (SSRIs) change the synaptic availability of serotonin. They can influence many forms of cognition, including learning [9–11], social perception [8], and decision-making [12]. They also alter central nervous system plasticity [13]. Given these central and higher-order cognitive effects of SSRI, we hypothesized that SSRIs may also alter higher-order processing of interoceptive signals.

There is good reason to expect a relationship between serotonin and interoception. Interoception communicates the state of homeostasis to influence cognition and emotion, contributes to the experience of reward and punishment, and is putatively susceptible to the regulatory orchestration of neural signals by other systems to determine its influence [14]. The serotonin system is also strongly linked to homeostasis-regulating processes including digestion, temperature, respiration, bladder control, and stress [15], is similarly implicated in a variety of cognitive and behavioral control processes [16–18] including representation of both reward and punishment [19], and has regulatory effects on the influence of other systems [18,20]. Anatomically, a broad distribution of the serotonin system in the brain reflects its influence on other processes and serotonin nuclei in the brainstem are well positioned to regulate communication between the brain and body. Serotonergic antidepressants have already been shown to modulate the experience of pain [21,22]. In the other direction, visceral states can alter central serotonin availability, potentially by peripheral regulation of tryptophan metabolism [23]. There are also hints of a relationship between serotonin and the cardiac interoceptive domain. In healthy individuals, a reduced correlation between neural and cardiac response to surprise, for instance, occurs when the serotonin precursor, tryptophan, is depleted [24]. Moreover, patients with serotonin-linked social anxiety disorder [25] have been shown to have reduced 5-HT1A binding potential in the insula cortex, a key area for interoception in the brain [3,26,27].

A significant advance for interoception research was the recognition that different dimensions of interoception vary independently in human experiments and provide markers for pharmacological testing [28,29]. Here, we focus on three: interoceptive accuracy (the tendency to make correct judgements about internal sensations), confidence in the judgements, and interoceptive insight (the metacognitive evaluation of interoceptive accuracy i.e., correspondence of confidence to accuracy of judgments). High interoceptive insight can occur as high confidence when judgements that are based on interoception are accurate and low confidence when judgements are inaccurate.

The ability to metacognitively distinguish between accurate and inaccurate interoceptive judgements, such as by improved predictions of incoming interoceptive information’s precision or other mechanism, could be the basis on which one regulates influence of interoception on mental states [2,14,30]. If this system falters, interoceptive sensations could have too much or too little influence, theoretically resulting in problems such as heightened anxiety [4], blunted affect [31], inappropriate response to hunger [5], poor choices [32,33] and reactive aggression [20]. Improved ability to distinguish informative and noninformative interoceptive signals, in contrast, could promote better control over interoceptive influence on other systems, and efficient shifts to alternative information sources for forming beliefs and inferences.

The heart has rich and bidirectional connections and so cardiac interoception is commonly tested [34]. As mentioned, cardiac interoception has indicated to have a relationship with serotonin [24] and was the interoceptive domain available to our expertise and facilities. So cardiac interoception was tested in the present study.

We tested the causal link between acute serotonin changes and interoception, with the focus on interoceptive accuracy, confidence, and interoceptive insight. We used a within-subject, placebo-controlled, crossover study of healthy participants on and off the SSRI, citalopram. This design controlled for both individual differences and effects of repeated task performance. Citalopram was chosen for its specificity to serotonin (3,800 times the affinity to the norepinephrine transporter and 10,000 times the affinity to the dopamine transporter [35]), tolerability, and common use.

Interoceptive ability was quantified using a heartbeat discrimination task [28,36]. This requires a participant to attend to interoceptive sensations and report whether auditory tones are in or out of time with their heartbeat, and then self-rate confidence in that judgement. Interoceptive accuracy, confidence, and interoceptive insight were tested in the context of this task. Effects on interoceptive insight were measured as the overall ability for confidence to distinguish between correct and incorrect judgements, and further broken down into changes of confidence for correct and incorrect judgements.

## Materials and Methods

### Experimental Design

This study used a double-blind, placebo-controlled within-subject, cross-over design. Participants underwent two test sessions, at least one week apart, under medical supervision. In one session they ingested 20mg citalopram (Cipramil) in a cellulose capsule, with extra space filled with microcrystalline cellulose which is an inactive ingredient of the citalopram tablet. In the other session, they received placebo (an identical capsule containing microcrystalline cellulose, which was also in the citalopram tablet). No-one who had contact with participants was aware of the treatment order, which was pseudo-randomized, balanced for sex, and coded by a researcher who was not present during testing. Capsules were manufactured according to good manufacturing practice [37].

### Participants

On a separate occasion, prior to testing, prospective participants undertook a screening session with a health questionnaire, heart rate and blood pressure monitoring by a medical doctor, and a structured clinical interview to determine any undiagnosed psychiatric conditions (Mini International Neuropsychiatric Interview; MINI [38]).

Exclusion criteria included: age under 18 or over 35 years; history of psychiatric disorder (including anxiety disorder, depression, eating disorder, psychosis and substance abuse disorder); presence of significant ongoing medical condition (including migraine, diabetes, epilepsy, glaucoma and hypertension); pregnancy or breastfeeding; currently taking any medication (excluding oral contraceptive pill); first-degree family history of bipolar disorder; Mini International Neuropsychiatric Interview (MINI) indication of: major depressive episode, manic episode, panic disorder, social phobia, OCD, PTSD, alcohol dependence, substance dependence, mood disorder with psychotic features, psychotic disorder, anorexia nervosa, bulimia nervosa, generalized anxiety disorder, or antisocial personality disorder. Participants were also instructed to abstain from alcohol or caffeine in the preceding 12 hours before the start of test sessions.

Fifty-one participants were recruited. Each participant was asked if they could feel their own pulse in their finger with the apparatus in place. Three were excluded for feeling their pulse in their finger against the apparatus and one for technical errors preventing data collection. Forty-seven participants were successfully tested (mean age 23 (*SD* = 3.9), 31 females, mean weight 64 kg (*SD* = 10.9). Given weights of recruited participants, the average citalopram dose was 0.34 mg/kg (*SD* = .05). Testing was conducted in two separate locations (test cubicles) and no difference of result was observed between locations. This study received ethical approval from the University of Sussex Sciences & Technology Cross-Schools Research Ethics Committee (ER/JL332/3, ER/JL332/9). Participants gave informed written consent.

### Procedure

Interoception was measured as part of a battery of tasks, including information sampling, visual metacognition (see supplemental material), and social decision-making. Behavioral testing was timed to begin at 3 hours after administration, corresponding to estimated peak plasma levels [39].

Citalopram can exhibit side effects (typically mild at the dose used here) including nausea, headache, and dizziness [40]. Visual analogue scales (VAS; from 0-100) assessed the presence of these three somatic effects (nausea, headache, dizziness). Additionally, five emotion/arousal related effects were assessed with VAS scales between pairs of antonyms: alert−drowsy, stimulated−sedated, restless−peaceful, irritable−good-humored, anxious−calm. Each measure was recorded three times: immediately following dosing, at the start behavioral testing and at the end. Mean scores for the two testing times were used in analyses, with paired t-tests to analyze whether significant differences occurred between citalopram and placebo sessions. Cardiac measures of heart rate (HR) and heart rate variability (the standard deviation of HR across intervals) were calculated at baseline and test time. Citalopram has been reliably shown to not affect blood pressure without interaction with other drugs [41,42] so blood pressure was not a dependent measure.

### Task

Participants were connected to a fingertip pulse oximeter to monitor cardiac events (Xpod with 8000SM sensor, Nonin Medical Inc., Minnesota, USA). The task was run in MATLAB (version 2018a, MathWorks) using a variant originally developed in [43].

The interoception task [28,44] is a two-alternative forced choice task, often called the heartbeat discrimination task. Participants were instructed beforehand that the computer would play a set of tones that would be in or out of sync with their heartbeat. During each trial, their heartbeat was measured in real-time, while a computer played a set of ten tones at either ∼250ms or ∼550ms after the R-wave [45]. These timings correspond respectively to judgements of maximum and minimum simultaneity (i.e., synchronous or delayed) between stimulus presentation and heartbeat [46] (see limitations section of discussion for consideration of other variable interval approaches). Following each trial, the participant was directed to respond to whether the tones were in or out of time with their heartbeats, and how confident they were in that answer using a VAS scale ranging from ‘total guess’ to ‘complete confidence’ on a scale of 1 to 10. Synchronous and delayed trials were presented in a pseudorandomized order.

Twenty trials (10 synchronous and 10 delayed) were completed in each session. Performance on 20 trials has been shown to correlate at *r* = .7 with performance of 100 trials [47] or *r* = .85 with performance on 50 [48], within subjects, and this number was chosen based on findings of prior literature at the time of design [28]. Since all comparisons were within-subject we minimized issues arising from individual differences in the noise associated with using 20 trials (between-subject correlations, for example, would require more trials [47]).

### Analysis

Accuracy scores were calculated by taking the mean number of correct responses for the session and dividing by the number of trials, resulting in a proportion correct. Confidence scores were computed as the mean of the trial-wise confidence VAS measure. Interoceptive insight was calculated by comparing confidence in correct choices and confidence in incorrect choices between drug and placebo conditions.

The effects on correct and incurred response confidence can be summarized as the area under the receiver-operating characteristic curve (AUC, MATLAB v R2020a) [49,50], measuring the correspondence of confidence to accuracy, independent from and unbiased by individual differences of confidence. Note AUC curves of accuracy (choice predicting correctness) provide the same information as proportion correct.

We used repeated-measures GLMs to model within-subject differences between drug and placebo repeated measures ANOVA (JASP v 0.14). Effects or near significant effects of citalopram, nausea, heartrate, and anxiety were present. So, a second repeated-measures GLM was conducted for each measure that included, change of heartrate, change of nausea, and change of anxiety as covariates. There was no interaction effect with or moderation by gender or order and inclusion of these factors did not change results, so these were not included in the reported statistics.

Further testing of the influence of nausea, anxiety, and heart rate was completed by mediation analysis (Fig S2, which includes the covariate analysis above as the direct effect [51]) and analysis of the main effects using abridged datasets in which heart rate, nausea and anxiety changes are very similar between placebo and drug conditions (see Supplemental Material).

## Results

Baseline performances on both tasks were consistent with previous studies (Table 1) [28]. Citalopram reduced heart rate (beats per minute (bpm): *t*(46) = 3.9, *p* = .01, mean diff *ΔM* = 4.0), increased nausea (on 100-point scale: *t*(46) = 2.9, *p* = .01, *ΔM* = 4.7) compared with placebo, and on trend increased anxiety (on VAS scale of calm to anxious (scale 0-100), *t*(46) = −1.8, *p* = .09, *ΔM* = 4.3. Citalopram did not change heart rate variability or any other measured physiological or subjective state (see Table S1).

**Table 1.** Effects of Citalopram on Interoception Task Performance

| | Mean (SD) | | $F(1,46)$ | $p$ | $\eta^2$ | with change of<br>HR, nausea, &<br>anxiety<br>covariates | |
| --- | --- | --- | --- | --- | --- | --- | --- |
| | Placebo | Citalopram | | | | $F(1,42)$ | $p$ |
| interoceptive accuracy | 0.56 (0.13) | 0.58 (0.16) | 0.57 | .45 | .01 | 0.10 | .75 |
| confidence | 5.03 (1.85) | 5.47 (1.48) | 3.60 | .06 | .07 | 3.05 | .09 |
| interoceptive insight | 0.50 (0.18) | 0.58 (0.17) | 6.51 | .01 | .12 | 5.09 | .03 |

Interoceptive accuracy was above 50% in both conditions (placebo *t*(46) = 3.2, *p* = .002, citalopram *t*(46) = 3.6, *p* = .001) but unchanged by citalopram (Table 1). Citalopram increased interoceptive insight with and without covariates of change to heartrate, nausea, and anxiety (Table 1, Fig 1, Fig S1). Approximately two-thirds of individuals showed this effect. Further investigation showed the effect on interoceptive insight was driven by increases of confidence for correct interoceptive judgements. There was an interaction between the effect of citalopram and whether the judgement was correct (*F*(1,46) = 6.91 *p* = .01; with covariates *F*(1,42) = 3.55, *p* = 0.07). Confidence for correct judgements was higher on citalopram (*F*(1,46) = 6.75, *p* = .01; with covariates *F*(1,42) = 5.32, *p* = .02) (Fig 1C). Confidence for incorrect judgements did not change (*ps* > .25) (Fig 1C). There was no interaction effect between the effect of citalopram and (i) change of nausea (F(1,43=.85, *p* =.36), (ii) heart rate (F(1,43 = 001, *p* = .98), or (iii) anxiety (F(1,43=.07, *p* = .79) on interoceptive insight.

**Fig 1.**
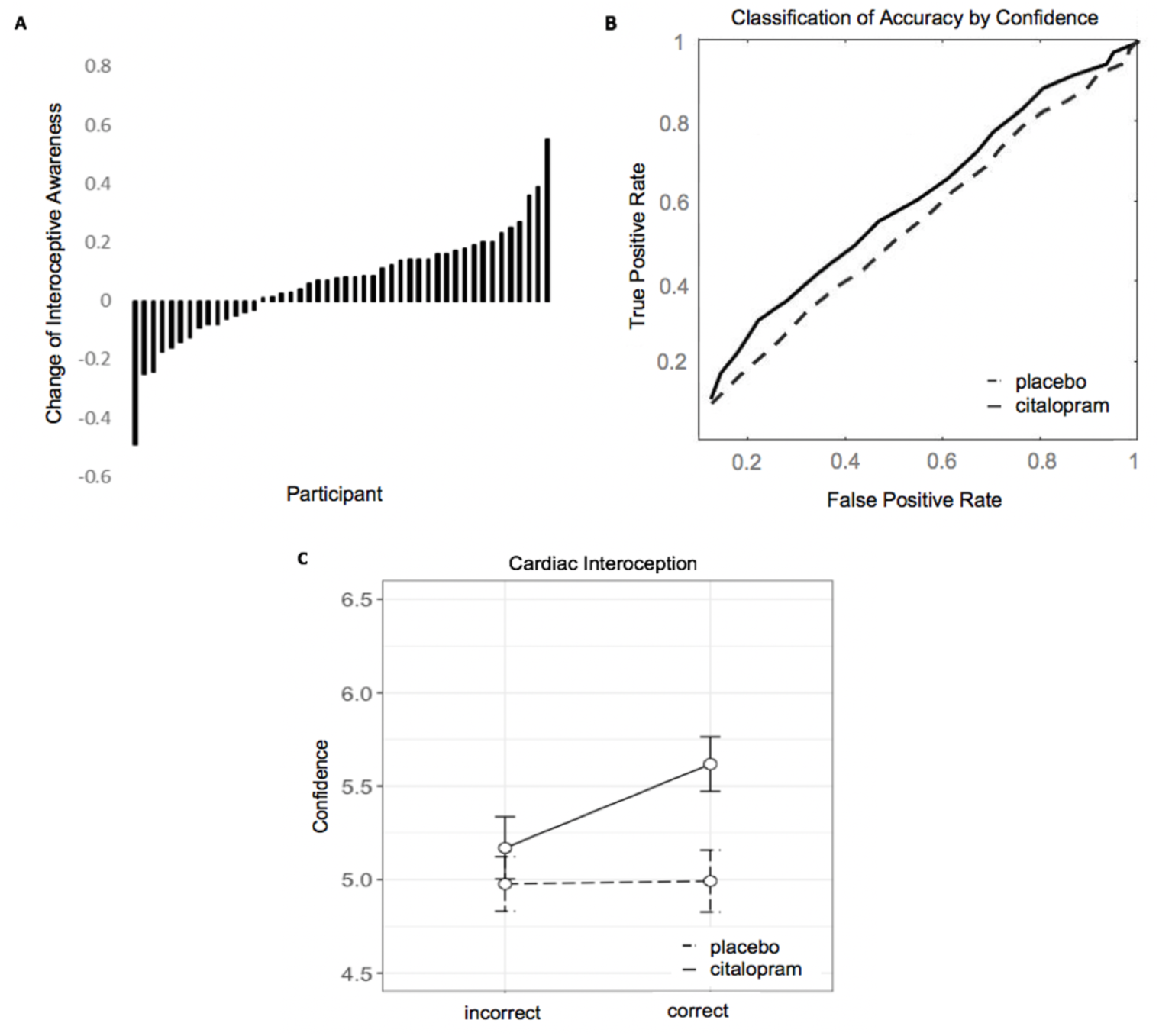
Effect of citalopram on interoceptive insight. **A**. Change in interoceptive insight by each participant. **B**. Receiver operating characteristic curve representing confidence classification of interoceptive accuracy. **C**. Confidence for correct and incorrect judgements on scale from 0 to 10. Error bars are within-subject standard error.

The independence of interoceptive insight from subjective and physiological effects of the drug was confirmed by follow up tests. We found near-zero correlation between interoceptive insight effects and changes of all physiological and subjective measures (Table S2), other than nausea which had a low non-significant correlation (*r*(45) =.14, *p* =.36). In restricted datasets with no difference between drug conditions on heart rate, anxiety, or nausea between drug conditions the effect of citalopram on interoceptive insight remained (see supplemental materials). The effect of citalopram on interoceptive insight was not mediated by citalopram effects on accuracy, confidence, or any other measured effect of the drug (Fig S2, Table S3).

In a separate experiment on the same participants, we investigated the effect of citalopram on a visual metacognition task (see supplemental material). Insight on both tasks correlated on placebo, but this relationship fell away on citalopram due to the greater effect of citalopram on interoceptive insight. Adding the change of visual metacognition as a covariate to the original analysis of citalopram’s effect on interoceptive insight, including all covariates, did not reveal a relationship between effects on the two tasks.

## Discussion

A single dose of the SSRI citalopram enhanced insight into the likelihood than an inference based on interoceptive information would be correct. This relationship was independent of citalopram effects on heartrate or self-reported subjective states. This demonstrated that an acute serotoninergic change is sufficient to change interoceptive metacognition. Below, we consider potential mechanisms, implications other cognition and serotonin function, limitations, and implications for clinical science.

The neurocognitive mechanism of citalopram’s effect on interoceptive insight could depend on the method or methods by which an individual determines accuracy. For example, if a particular allocation of attention (between sound, cardiac interoception, and other information) increases the likelihood of a correct choice, then improved retrospective awareness of attentional allocation would improve insight into performance [52]. Alternatively, the mechanism could be that enhanced serotonin transmission increases insight into the quality of interoceptive information used to make the choice [53]. Predictive coding models explain subjective feelings and allostasis as outputs of interactions between top-down predictions about the states of the body and environment, and new, bottom-up sensations [14,30,54,55]. Layers of processing are implemented by similarly hierarchical neural networks, through which top-down predictions suppress or explain away signals encoded at lower levels. This leaves remaining signals to be broadcast forward to adjust higher-order predictions and drive allostasis (physiological, cognitive, and behavioural responses to changes within the body) [14,30,55]. With respect to the current findings, the top-down predictions posited to include ‘confidence’ in the lower-level prediction errors, which in turn, can be used to mediate post-synaptic gain [56]. This gain can ease the propagation of bottom-up information to influence higher levels of the network [57]. Clinical disorders associated with altered interoception have been considered within this predictive processing framework. Anxiety disorders, for instance, are proposed to arise from suboptimal interactions between lower and higher-level processes, whereby overreliance on incorrect prior expectations rather than useful lower-level inputs lead to constant surprise from new information and a perpetual drive for correction [4]. Improved interoceptive insight could help remediate such maladaptive cycles through increased reliance on reliable interoceptive inputs [53,58]. Given its regulatory effects on information flow in other systems [18,20], serotonin could theoretically mediate access to information about the precision of upward interoceptive signals. Precision itself may be encoded by other systems with which serotonin interacts (e.g., dopamine). Regardless of mechanism, if confidence in reliable interoceptive information is increased on citalopram, then the individual is better equipped to efficiently regulate the use of interoceptive signals, making consequent beliefs and feelings more accurate reflections of reality.

The interoception task is challenging. Accuracy was above chance in both treatment conditions, but only a minority performed above chance in both sessions. The challenging nature of the task provides a poor fit for signal detection approaches to metacognition analysis that assume that metacognitive insight arises from the same information and processes as the perceptual decision, necessitating above chance accuracy to interpret above chance metacognition [59]. However, decision-making and metacognition can have different inferential goals and be represented by different anatomy (see [60]). They can therefore be receptive to distinct types of information or be determined by different processing of the same information. These differences can lead to situations of high metacognitive sensitivity despite low first-order accuracy [61], and indeed, high metacognitive accuracy (‘blindsight’) has been demonstrated at robustly chance performance [60]. In the present study, the effect of citalopram on interoceptive insight fits with top-down regulatory roles of serotonin, whereby citalopram could alter top-down insight into the choice process, without necessarily altering the choice itself. This specificity provides a pharmacological dissociation between choice and insight, with new implications for serotonin’s role in perception and consciousness.

Targeted blockade of serotonin transporters has indirect effects on other systems [62]. So, while a shift of serotonin is sufficient to change interoceptive insight, the mechanism could also involve other neurotransmitters. Moreover, the effect could stem from either activation or inhibition of serotonin transmission. Acute SSRI treatment can cause reductions or increases in serotonin transmission due to activation of 5-HT1A auto-receptors, with variation of this across brain regions [63]. Moreover, effects of chronic citalopram treatment on interoception are not yet known [8].

Finally, any effect on discrimination task performance may result from enhanced insight into either processing interoceptive information or interoceptive-exteroceptive signal integration.

### Effects on other cognition

Accurate higher order representations of the ability to make good interoceptive inferences could enable better top-down regulation, consciously or unconsciously, of interoceptive influence on other cognitive and emotional processes. In turn, this could result in mental states that are better reflections of reality (2,64). The onward effects of these changes in interoceptive insight on cognition and mood are important and exciting avenues for future research.

In a separate experiment, we did not observe an effect of citalopram in the same participants for visual metacognition (Table S4). Correspondingly, the effect on interoceptive insight was larger than a metacognitive effect on the visual task. Reflecting this, a significant correlation of metacognitive insight measures on the two tasks on placebo falls away on citalopram. The design differences between the tasks do not make them directly comparable (i.e., number of trials and difference of modalities (visual vs interoceptive/auditory), which is why this analysis is supplemental to our main findings. Taken with caution, however, this may be the initial sign of the selectivity of serotonin’s effect on interoceptive insight rather than on metacognition in general.

Further work is needed to show whether the serotonergic effect on cardiac interoception generalizes to other interoceptive domains such as respiratory or gastric sensations. To this issue, cardiac and respiratory interoceptive insight have been positively correlated in prior work [64].

### Implications for serotonin function

The pursuit of describing a unifying function for serotonin has been long and informative but not entirely forthcoming [65]. This finding of a causal link between serotonin and interoception, together with involvement of both interoception and serotonin in aversive and rewarding signal processing [19,66], impulsive aggression [20,67], startle [68,69] and perception of threat [8,70] sets the stage for more overlap of mechanism across a diversity of serotonin effects on affective cognition than previously thought.

### Clinical Implications

The effect of citalopram on interoceptive insight provides a new framework for the study of serotonin in disorders associated with altered interoception. SSRI treatments and successful interoceptive therapies in depression and anxiety [71] may be found to overlap in mechanism as they ameliorate blunted affect [4] or break maladaptive cognitive cycles [72]. Perceptual biases that appear long before clinical effects of SSRIs are observed and thought to predict clinical outcomes [8,10] may be discovered to have interoceptive foundations. Anxiogenic effects arising near the onset of SSRI treatment [73] may be due interoceptive inputs suddenly being processed in a new way. If these hypotheses are supported in future studies, clinical effectiveness of SSRI treatment might be predicted by early interoceptive effects.

### Limitations

Research published after this project was designed noted that high numbers of trials (∼40-60) are recommended to attain exceptionally high reliability on interoceptive accuracy, measured as an exceptionally high correlation (r > 0.9) with and accuracy levels measured with 100 trials [47]. The correlation of accuracy on 20 trials with the accuracy on 100 less ideal but still reasonable (r = 0.7). So, the lack of citalopram effect on interoceptive accuracy (first order performance on the task), could be interpreted with the caution that effects may differ when more trials are used. The within-subject design used in this study removes issues highlighted for between-subject comparisons and correlations. Critically, the citalopram effect on interoceptive insight is not reliant on a specific level of accuracy, but rather awareness of when accuracy was high. Also related to between-subject comparisons, it has been recommended to use multiple intervals between the R-wave and tones to account for individual differences in what is perceived to be in sync and out of sync with the heart when assigning a particular accuracy level to an individual [74]. Like most prior studies, we used a single interval for all participants. The within-subject design of this study mitigates problems of a single interval because any variance of subjective sense of in and out of sync beats would be the same on and off citalopram. Yet, if there is substantial between-subject variance of intervals associated with being in sync with one’s heartbeat, then more information on the precise nature of the effect would be gained future studies using variable intervals.

The difficulty (low first order accuracy) of the task could mean that effects of citalopram on interoceptive insight could theoretically and intriguingly be limited to situations of high uncertainty, requiring further study across an array of task difficulties to fully understand applicable contexts.

### Summary

With this study, we find that serotonin can alter interoceptive metacognitive insight. In doing so, we provide evidence for serotonin activity a potential moderator of the ability to make reliable inferences from sensations within the body.

## Acknowledgments

We would like to thank Dr Stephen Fleming for providing feedback and Dr. Lina Skora for advice and comments on earlier drafts of this report.

## Author contributions

JL and DC-M designed the study in consultation with HC and SG. JL, CH, and KA collected data. GM conducted medical screening and medical support during data collection. DC-M and JL conducted analyses and wrote the manuscript with edits and approval from all authors.

## Data and materials availability

All data, code, and materials used in the analysis are available any researcher for purposes of reproducing or extending the analysis. To download the data associated with this manuscript go to https://figshare.com/s/7154cc6d436e54154c37.

## Supplementary Materials for

### Supplementary Results

**Fig S1.**
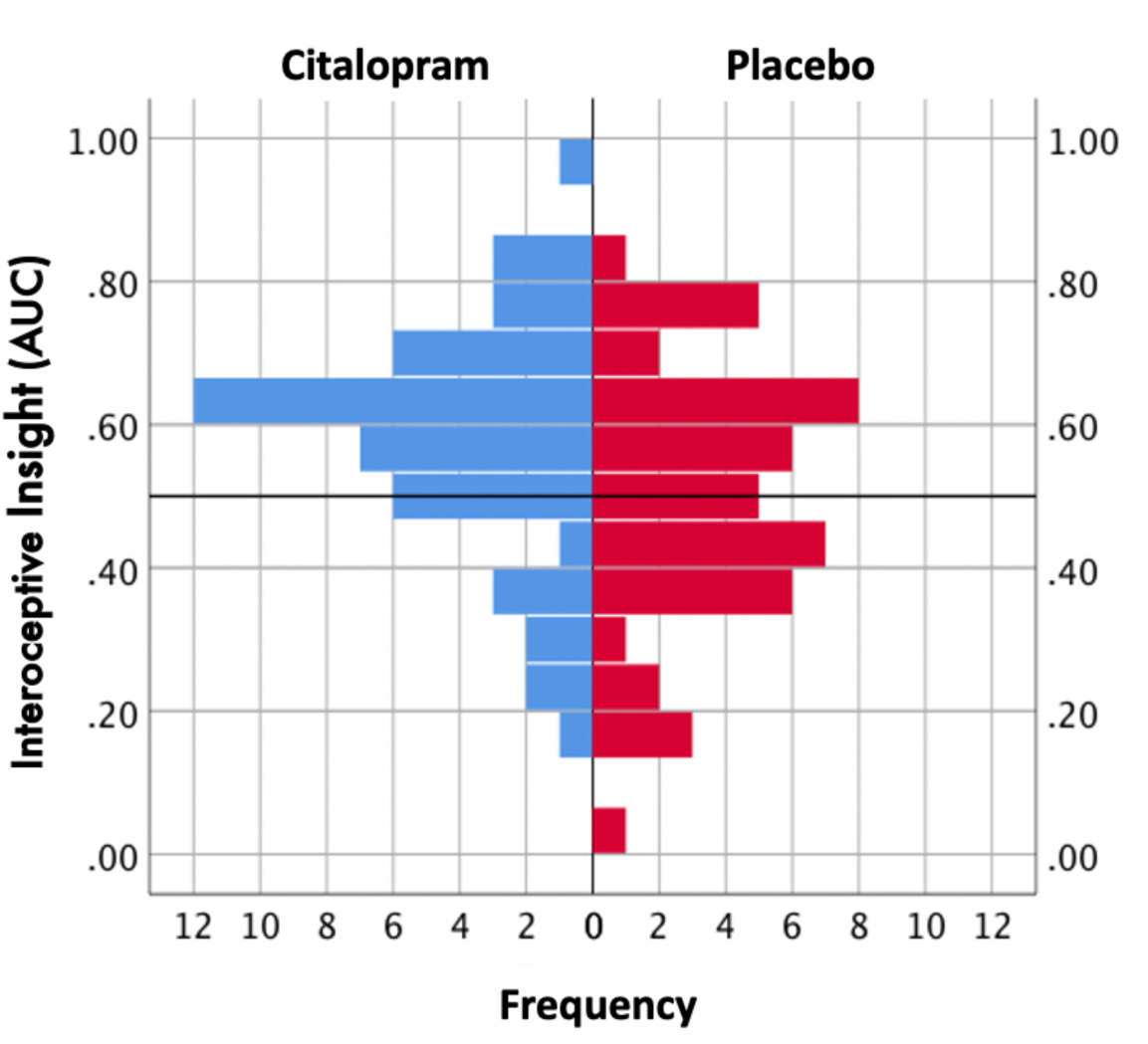
Distribution of Interoceptive Insight in both Conditions.

#### Somatic and psychological effects

Table S1 shows the difference between scores in drug and placebo conditions on subjective ratings at test times. There were differences in the sample on the subjective nausea rating, corresponding to a mean 4.7 difference on a 100-point scale. There was also a drop of heart rate on citalopram.

**Table S1:** Mean VAS score (SD) at test time and contrasts between drug conditions. † full sample, ‡ restricted sample, * p < .05, ** p < .01

|  | scale | placebo | citalopram | *t*(46) | *p* |
| --- | --- | --- | --- | --- | --- |
| FULL SAMPLE | nausea | 4.47 (7.29) | 9.2 (11.8) | -2.93 | .005** |
|  | headache | 12.3 (17.3) | 12.2 (17.6) | 0.05 | .96 |
|  | dizziness | 9.4 (13.0) | 11.0 (12.1) | -1.22 | .23 |
|  | alert – drowsy | 44.0 (18.2) | 47.8 (19.5) | -1.27 | .21 |
|  | stimulated – sedated | 46.4 (15.6) | 47.7 (14.9) | -0.54 | .59 |
|  | restless – peaceful | 64.3 (18.6) | 61.2 (18.7) | 1.02 | .31 |
|  | irritable – good-humoured | 63.9 (17.7) | 64.8 (16.1) | -0.49 | .62 |
|  | anxious – calm | 71.9 (15.9) | 67.6 (15.7) | 1.75 | .086 |

In addition to reporting results with change of heart rate, anxiety and nausea as covariates in the main analysis, we did further analysis to confirm the absence of influence of these factors on our findings. First, we tested for correlations of each change with the effect of citalopram on interoceptive insight (Table S2). No changes in cardiac or self-report variables between drug and placebo were significantly related to interoceptive insight. Next, we conducted a second analysis on restricted datasets, whereby cases were removed until a statistical comparison between citalopram and placebo exceeded *p* > 0.8. If data is restricted (N=31) to no difference of heart rate (F(1,30)=.01, *p* = .92, *ΔM* − 0.08, SD 4.4 bpm reduction) between drug conditions, the effect of citalopram on interoceptive insight remains (*F*(1,30) = 5.14 *p* = .03). If data is restricted (N=37) to no difference on nausea (F(1,36)=.027, *p* = .87, *ΔM* = .13 SD 4.98), the citalopram effect on interoceptive insight remains (*F*(1,36) = 5.0, *p* = .03). If data is restricted (N=42) to no difference on anxiety (F(1,41) = .03, *p* =.87, *ΔM* = .33 SD 12.79), the effect of citalopram on interoceptive insight remains (F(1,41) = 4.2, *p* =.046).

**Table S2:** Correlations between drug-placebo changes in cardiac/self-report variables and interoceptive insight

| Change of: | interoceptive insight |  |
| --- | --- | --- |
|  | *r*(45) | p |
| heart rate | .01 | .97 |
| heart rate variability | -.01 | .97 |
| nausea | -.14 | .36 |
| headache | .00 | .98 |
| dizziness | .04 | .82 |
| alert–drowsy | .02 | .90 |
| stimulated – sedated | -.08 | .59 |
| restless – peaceful | .01 | .95 |
| irritable – good-humoured | -.01 | .94 |
| anxious – calm | .0 | .88 |

Finally, we conducted a mediation analysis (Fig S2) to assess the possibility of indirect citalopram effects on interoceptive insight via other measured factors. We ran the mediation analyses using a within-subjects approach (MEMORE v2.1), which also includes both changes and average values of the mediator across conditions in the model [1]. This was conducted for all potential mediators showing differences in the mediating variable between drug and placebo i.e., path B, or correlations between drug-placebo differences and task variables (path C). Results are reported in Table S3. No significant mediations were found.

**Fig S2:**
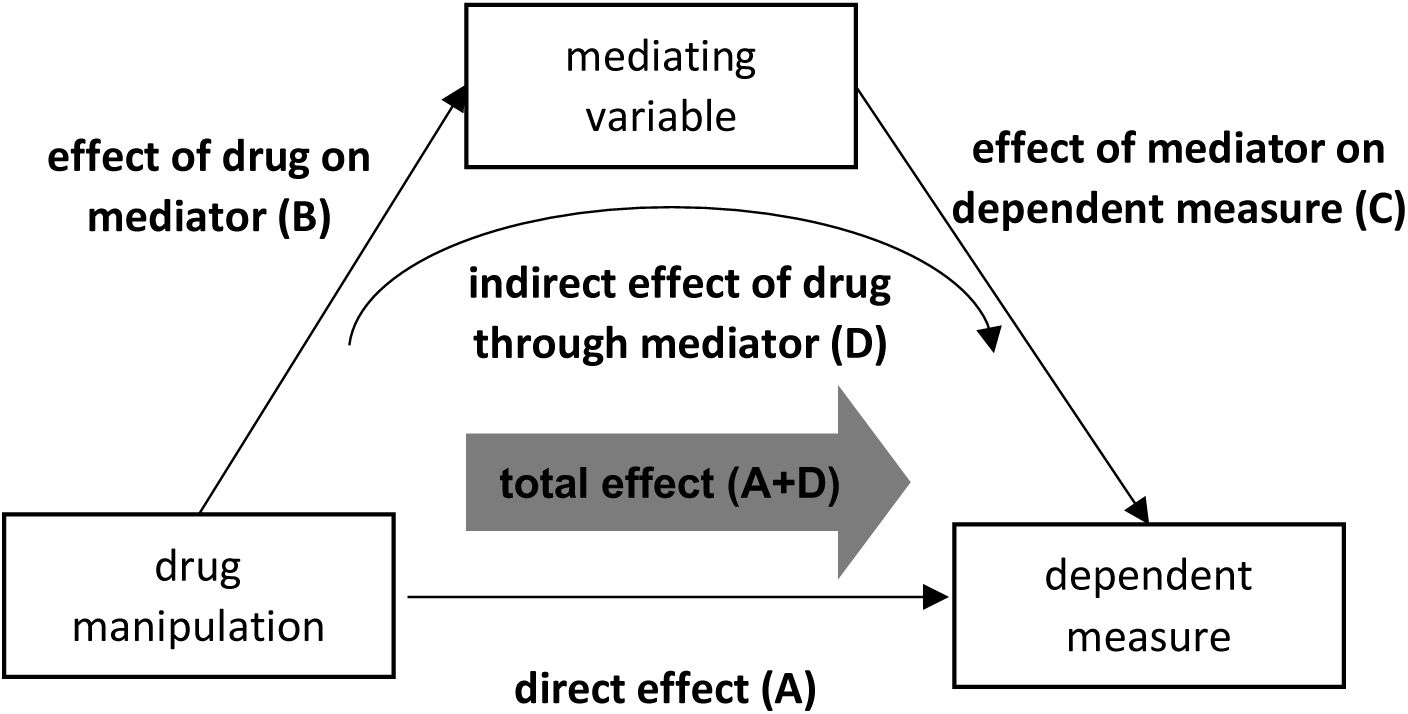
Mediation analysis approach.

**Table S3:** mediation analyses on interoceptive insight. HR – heart rate, † in units of insight scores, ‡ in units of the mediator. If *b*ootstrap confidence intervals overlap zero, *indirect* effect is determined to be non-significant.

|  | Mediator(s) |  |  |  |
| --- | --- | --- | --- | --- |
|  | HR | nausea | anxiety | ALL 3 |
| direct citalopram effect on interoceptive insight † | 0.08 | 0.07 | 0.07 | 0.08 |
| *t*-stat | 2.27 | 2.33 | 2.45 | 2.33 |
| *p* | **.028** | **.024** | **.018** | **.024** |
| effect of citalopram on mediator ‡ | -3.99 | 4.71 | 4.29 |  |
| *t*-stat | -3.91 | 2.93 | 1.75 |  |
| *p* | **<.001** | **.005** | .086 |  |
| effect of mediator on interoceptive insight † | 0.0013 | <0.0001 | .0002 |  |
| *t*-stat | 0.27 | 0.01 | .10 |  |
| *p* | .77 | .99 | .86 |  |
| indirect effect of citalopram on interoceptive insight through mediator (path D) † | -0.005 | -0.0002 | -.001 |  |
| Bootstrap Lower / Upper CI | -.05/.03 | -.04/.02 | -.01/.01 |  |

### Supplementary Experiment

The same participants a visual metacognitive Insight task (VMI) after the interoception task.

#### Methods

##### VMI task

The visual task was taken from [2]. Participants were shown circles containing dots and instructed to indicate which contained more. Following each trial, they were asked to indicate their confidence in the previous response on a Likert scale. 200 trials were conducted in 8 blocks, with a self-timed rest every 25 trials. The difficulty was staircased over the course of the task, with the difference in numbers of dots (Δd), adjusted to target a mean rate of correct answers of 70%, to keep a consistent level of difficulty between participants. One randomly selected circle always held 50 dots. After two consecutive correct responses, Δd was decreased by one dot; after one incorrect response, Δd was increased by one dot.

We calculated Δd corresponding to the number of dots differing between the two circles necessary to maintain 70% accuracy as a measure of performance. Staircasing was successful: mean accuracy was .71 (*SD* = .02) on placebo and .72 (*SD* = .02) on citalopram. Confidence scores were recorded on each trial. Using performance and confidence ratings (1 to 6), we calculated visual metacognitive insight (VMI) as AUC, corresponding to the interoceptive insight measure. This common measure allowed a direct comparison of citalopram effects on cardiac interoception and visual exteroception. We could then look specifically for changes of confidence on correct and incorrect judgements.

We also calculated visual metacognitive sensitivity (meta d’) and efficiency (meta d’ / d’) for reference to previous research [3,4]. Correlation of VMI and meta d’ in the placebo condition was *r* = .92, and correlation between VMI and meta d’/d’ was *r* = .86. We used GLMs including order, nausea, heart rate and anxiety as regressors of no interest to model within-subject differences between drug and placebo. We also examined the effects on confidence independently in correct and incorrect choices.

For comparison of VMI with interoceptive insight controlling for the difference in accuracy and number of trials between tasks, we completed a supplementary analysis on VMI. 1000 random draws of 20 samples each were taken from each participant separately for drug and placebo sessions, weighted to include correct and incorrect trials at the same proportion as that person’s interoception task accuracy, in the same session. Computations on insight for each sample were made and the average of these used in statistical comparison with interoceptive insight at the same level of accuracy.

To test if citalopram effects on interoception were over and above a general effect on metacognition, we performed a post hoc analysis of interoceptive insight with the addition of change of visual metacognition as a covariate in our original analyses.

#### Results

Interoceptive insight correlated with its equivalent measure in visual perception (VMI) (*r*(45) = .35, *p* = .02) in the placebo condition indicating some consistency between measures, but this relationship disappeared on citalopram (*p* = .85).

Given an observation of near significant interaction with order of treatment for some measures of visual metacognition, order was included as a between subject factor in all models in addition to change of nausea, heart rate and anxiety.

A comparison of citalopram effects between interoceptive insight and VMI demonstrated a significant drug × task interaction (*F*(1,45) = 6.56, *p* = .014), with a greater effect on interoception. There was no effect of citalopram on VMI (Table S4). This remains the case for subsets of visual perception data matching interoceptive task accuracy (20 visual task trials with same mean accuracy as interoceptive task performance).

Inclusion of visual metacognition changes in the original interoceptive insight analysis (in addition to anxiety, nausea and heart rate) as a covariate did not predict or change the effect of citalopram on interoceptive insight (F(1,42)=5.78, *p* = .02). Change of visual metacognition did not predict the citalopram effects on interoceptive insight (F1,42) = 0.22, *p* = .64). There was no interaction effect between effects of citalopram on VMI changes and interoceptive insight (F(1,42)=1.60, *p* = 0.21).

**Table S4:** Repeated measures ANOVA conditional main effect for all exteroception (visual) metacognition task. Full factorial models include order as a between subject factor, change of nausea, change of heartbeat, and change of anxiety, interactions of each covariate and factor with drug condition.

| | placebo | citalopram | conditional<br>main effect<br>$F(1,45)$ | $p$ |
| --- | --- | --- | --- | --- |
| $\Delta d$ | 5.49 | 5.74 | 0.00 | .99 |
| confidence (1-10 scale) | 5.84 | 5.66 | 0.50 | .48 |
| confidence correct<br>choices | 6.19 | 5.99 | 0.91 | .35 |
| confidence incorrect<br>choices | 5.00 | 4.79 | .01 | .94 |
| VMI (AUC) | 0.65 | 0.65 | .50 | .49 |
| meta $d'$ | 0.96 | 0.96 | 0.48 | .49 |
| meta $d' / d'$ | 0.79 | 0.77 | 1.19 | .28 |

## References

1. Cameron OG. Visceral Sensory Neuroscience. Oxford: Oxford University Press; 2002.

2. Seth AK. Interoceptive inference, emotion, and the embodied self. Trends Cogn Sci (Regul Ed). 2013;17:565–573.

3. Craig A. How do you feel? Interoception: the sense of the physiological condition of the body. Nat Rev Neurosci. 2002;3:655–666.

4. Paulus MP, Feinstein JS, Khalsa SS. An Active Inference Approach to Interoceptive Psychopathology. Annual Review of Clinical Psychology. 2019;15:97–122.

5. Herbert B, Pollatos O. The relevance of interoception for eating behavior and eating disorders. The Interoceptive Mind, Oxford: Oxford University Press; 2019. p. 165–186.

6. Avery JA, Drevets WC, Moseman SE, Bodurka J, Barcalow JC, Simmons WK. Major depressive disorder is associated with abnormal interoceptive activity and functional connectivity in the insula. Biological Psychiatry. 2014;76:258–266.

7. Ehlers A. Interoception and panic disorder. Advances in Behaviour Research and Therapy. 1993;15:3–21.

8. Harmer CJ, Cowen PJ. ‘It’s the way that you look at it’—a cognitive neuropsychological account of SSRI action in depression. Philosophical Transactions of the Royal Society B: Biological Sciences. 2013;368:20120407.

9. Chamberlain SR, Muller U, Blackwell AD, Clark L, Robbins TW, Sahakian BJ. Neurochemical modulation of response inhibition and probabilistic learning in humans. Science. 2006;311:861–863.

10. Michely J, Eldar E, Martin IM, Dolan RJ. A mechanistic account of serotonin’s impact on mood. Nature Communications. 2020;11:2335.

11. Scholl J, Kolling N, Nelissen N, Browning M, Rushworth MFS, Harmer CJ. Beyond negative valence: 2-week administration of a serotonergic antidepressant enhances both reward and effort learning signals. PLoS Biol. 2017;15.

12. Crockett MJ, Clark L, Hauser MD, Robbins TW. Serotonin selectively influences moral judgment and behavior through effects on harm aversion. Proc Natl Acad Sci U S A. 2010;107:17433–17438.

13. Castrén E, Rantamäki T. The role of BDNF and its receptors in depression and antidepressant drug action: Reactivation of developmental plasticity. Developmental Neurobiology. 2010;70:289–297.

14. Owens AP, Allen M, Ondobaka S, Friston KJ. Interoceptive inference: From computational neuroscience to clinic. Neuroscience & Biobehavioral Reviews. 2018;90:174–183.

15. Berger M, Gray JA, Roth BL. The Expanded Biology of Serotonin. Annu Rev Med. 2009;60:355–366.

16. Cools R, Nakamura K, Daw ND. Serotonin and Dopamine: Unifying Affective, Activational, and Decision Functions. Neuropsychopharmacology. 2011;36:98–113.

17. Deakin JF, Graeff FG. 5-HT and mechanisms of defence. J Psychopharmacol (Oxford). 1991;5:305–315.

18. Dayan P, Huys QJM. Serotonin in Affective Control. Annual Review of Neuroscience. 2009;32:95–126.

19. Cohen JY, Amoroso MW, Uchida N. Serotonergic neurons signal reward and punishment on multiple timescales. Elife. 2015;4:e06346.

20. Spoont MR. Modulatory role of serotonin in neural information processing: Implications for human psychopathology. Psychological Bulletin. 1992;112:330–350.

21. Micó JA, Ardid D, Berrocoso E, Eschalier A. Antidepressants and pain. Trends in Pharmacological Sciences. 2006;27:348–354.

22. Jann MW, Slade JH. Antidepressant Agents for the Treatment of Chronic Pain and Depression. Pharmacotherapy: The Journal of Human Pharmacology and Drug Therapy. 2007;27:1571–1587.

23. O’Mahony SM, Clarke G, Borre YE, Dinan TG, Cryan JF. Serotonin, tryptophan metabolism and the brain-gut-microbiome axis. Behavioural Brain Research. 2015;277:32–48.

24. Mueller EM, Evers EA, Wacker J, Van Der Veen F. Acute tryptophan depletion attenuates brain-heart coupling following external feedback. Front Hum Neurosci. 2012;6.

25. Frick A, Åhs F, Engman J, Jonasson M, Alaie I, Björkstrand J, et al. Serotonin Synthesis and Reuptake in Social Anxiety Disorder: A Positron Emission Tomography Study. JAMA Psychiatry. 2015;72:794.

26. Lanzenberger RR, Mitterhauser M, Spindelegger C, Wadsak W, Klein N, Mien L-K, et al. Reduced Serotonin-1A Receptor Binding in Social Anxiety Disorder. Biological Psychiatry. 2007;61:1081–1089.

27. Schulz SM. Neural correlates of heart-focused interoception: a functional magnetic resonance imaging meta-analysis. Philosophical Transactions of the Royal Society B: Biological Sciences. 2016;371:20160018.

28. Garfinkel SN, Seth AK, Barrett AB, Suzuki K, Critchley HD. Knowing your own heart: Distinguishing interoceptive accuracy from interoceptive awareness. Biological Psychology. 2015;104:65–74.

29. Khalsa SS, Adolphs R, Cameron OG, Critchley HD, Davenport PW, Feinstein JS, et al. Interoception and Mental Health: A Roadmap. Biological Psychiatry: Cognitive Neuroscience and Neuroimaging. 2018;3:501–513.

30. Allen M, Tsakiris M. The body as first prior: Interoceptive predictive processing and the primacy of self-models. Oxford University Press; 2018.

31. Paulus MP, Stein MB. Interoception in anxiety and depression. Brain Struct Funct. 2010;214:451–463.

32. Damasio AR. The somatic marker hypothesis and the possible functions of the prefrontal cortex. Philosophical Transactions of the Royal Society B: Biological Sciences. 1996;351:1413–1420.

33. Werner NS, Jung K, Duschek S, Schandry R. Enhanced cardiac perception is associated with benefits in decision-making. Psychophysiology. 2009;46:1123–1129.

34. Tsakiris M. The multisensory basis of the self: From body to identity to others. Quarterly Journal of Experimental Psychology. 2017;70:597–609.

35. Owens MJ, Knight DL, Nemeroff CB. Second-generation SSRIs: human monoamine transporter binding profile of escitalopram and R-fluoxetine. Biol Psychiatry. 2001;50:345–350.

36. Katkin E, Reed S, Deroo C. A methodological analysis of 3 techniques for the assessment of individual-differences in heartbeat detection. vol. 20, SOC PSYCHOPHYSIOL RES 1010 VERMONT AVE NW SUITE 1100, WASHINGTON, DC 20005; 1983. p. 452–452.

37. European Medicines Agency. Good manufacturing practice. European Medicines Agency. 2018. https://www.ema.europa.eu/en/human-regulatory/research-development/compliance/good-manufacturing-practice. Accessed 8 December 2020.

38. Sheehan DV, Lecrubier Y, Sheehan KH, Amorim P, Janavs J, Weiller E, et al. The Mini-International Neuropsychiatric Interview (M.I.N.I.): the development and validation of a structured diagnostic psychiatric interview for DSM-IV and ICD-10. The Journal of Clinical Psychiatry. 1998;59 Suppl 20:22-33;quiz 34-57.

39. Milne RJ, Goa KL. Citalopram. A review of its pharmacodynamic and pharmacokinetic properties, and therapeutic potential in depressive illness. Drugs. 1991;41:450–477.

40. Ekselius L, von Knorring L, Eberhard G. A double-blind multicenter trial comparing sertraline and citalopram in patients with major depression treated in general practice. Int Clin Psychopharmacol. 1997;12:323–331.

41. Zhong Z, Wang L, Wen X, Liu Y, Fan Y, Liu Z. A meta-analysis of effects of selective serotonin reuptake inhibitors on blood pressure in depression treatment: outcomes from placebo and serotonin and noradrenaline reuptake inhibitor controlled trials. Neuropsychiatr Dis Treat. 2017;13:2781–2796.

42. Watts SW, Morrison SF, Davis RP, Barman SM. Serotonin and blood pressure regulation. Pharmacol Rev. 2012;64:359–388.

43. Hart N, McGowan J, Minati L, Critchley HD. Emotional regulation and bodily sensation: interoceptive awareness is intact in borderline personality disorder. J Pers Disord. 2013;27:506–518.

44. Katkin ES, Morell MA, Goldband S, Bernstein GL, Wise JA. Individual Differences in Heartbeat Discrimination. Psychophysiology. 1982;19:160–166.

45. Payne RA, Symeonides CN, Webb DJ, Maxwell SRJ. Pulse transit time measured from the ECG: an unreliable marker of beat-to-beat blood pressure. J Appl Physiol. 2006;100:136–141.

46. Wiens S, Palmer SN. Quadratic trend analysis and heartbeat detection. Biological Psychology. 2001;58:159–175.

47. Kleckner IR, Wormwood JB, Simmons WK, Barrett1 LF, Quigley KS. Methodological Recommendations for a Heartbeat Detection-Based Measure of Interoceptive Sensitivity. Psychophysiology. 2015;52:1432–1440.

48. Jones GE, Jones KR, Rouse CH, Scott DM, Caldwell JA. The Effect of Body Position on the Perception of Cardiac Sensations: An Experiment and Theoretical Implications. Psychophysiology. 1987;24:300–311.

49. Green DM, Swets JA. Signal detection theory and psychophysics. Oxford, England: John Wiley; 1966.

50. Hajian-Tilaki K. Receiver Operating Characteristic (ROC) Curve Analysis for Medical Diagnostic Test Evaluation. Caspian J Intern Med. 2013;4:627–635.

51. Montoya AK, Hayes AF. Two-condition within-participant statistical mediation analysis: A path-analytic framework. Psychological Methods. 2017;22:6–27.

52. Kanai R, Walsh V, Tseng C. Subjective discriminability of invisibility: A framework for distinguishing perceptual and attentional failures of awareness. Consciousness and Cognition. 2010;19:1045–1057.

53. Yeung N, Summerfield C. Metacognition in human decision-making: confidence and error monitoring. Philosophical Transactions of the Royal Society B: Biological Sciences. 2012;367:1310–1321.

54. Seth AK. Interoceptive inference, emotion, and the embodied self. Trends in Cognitive Sciences. 2013;17:565–573.

55. Friston K. Predictive coding, precision and synchrony. Cognitive Neuroscience. 2012;3:238–239.

56. Abbott LF, Varela JA, Sen K, Nelson SB. Synaptic Depression and Cortical Gain Control. Science. 1997;275:221–224.

57. Feldman H, Friston K. Attention, Uncertainty, and Free-Energy. Frontiers in Human Neuroscience. 2010;4:215.

58. Spence ML, Dux PE, Arnold DH. Computations underlying confidence in visual perception. Journal of Experimental Psychology: Human Perception and Performance. 2016;42:671–682.

59. Barrett AB, Dienes Z, Seth AK. Measures of metacognition on signal-detection theoretic models. Psychological Methods. 2013;18:535–552.

60. Scott RB, Dienes Z, Barrett AB, Bor D, Seth AK. Blind Insight: Metacognitive Discrimination Despite Chance Task Performance. Psychol Sci. 2014;25:2199–2208.

61. Fleming SM, Daw ND. Self-Evaluation of Decision-Making: A General Bayesian Framework for Metacognitive Computation. Psychol Rev. 2017;124:91–114.

62. Zhou F-M, Liang Y, Salas R, Zhang L, Biasi MD, Dani JA. Corelease of Dopamine and Serotonin from Striatal Dopamine Terminals. Neuron. 2005;46:65–74.

63. Beyer CE, Cremers TIFH. Do selective serotonin reuptake inhibitors acutely increase frontal cortex levels of serotonin? European Journal of Pharmacology. 2008;580:350–354.

64. Garfinkel SN, Manassei MF, Hamilton-Fletcher G, In den Bosch Y, Critchley HD, Engels M. Interoceptive dimensions across cardiac and respiratory axes. Philosophical Transactions of the Royal Society B: Biological Sciences. 2016;371:20160014.

65. Dayan P, Huys Q. Serotonin’s many meanings elude simple theories. ELife. 2015;4.

66. Marshall AC, Gentsch A, Blum A-L, Broering C, Schütz-Bosbach S. I feel what I do: Relating interoceptive processes and reward-related behavior. NeuroImage. 2019;191:315–324.

67. Coccaro EF, Fanning JR, Phan KL, Lee R. Serotonin and impulsive aggression. CNS Spectr. 2015;20:295–302.

68. Grillon C, Levenson J, Pine DS. A single dose of the selective serotonin reuptake inhibitor citalopram exacerbates anxiety in humans: a fear-potentiated startle study. Neuropsychopharmacology. 2007;32:225–231.

69. Schulz A, Matthey JH, Vögele C, Schaan V, Schächinger H, Adler J, et al. Cardiac modulation of startle is altered in depersonalization-/derealization disorder: Evidence for impaired brainstem representation of baro-afferent neural traffic. Psychiatry Research. 2016;240:4–10.

70. Garfinkel SN, Critchley HD. Threat and the Body: How the Heart Supports Fear Processing. Trends in Cognitive Sciences. 2016;20:34–46.

71. Khoury NM, Lutz J, Schuman-Olivier Z. Interoception in Psychiatric Disorders: A Review of Randomized, Controlled Trials with Interoception-Based Interventions. Harv Rev Psychiatry. 2018;26:250–263.

72. Roy-Byrne PP, Craske MG, Stein MB. Panic disorder. The Lancet. 2006;368:1023–1032.

73. Taylor MJ, Freemantle N, Geddes JR, Bhagwagar Z. Early Onset of Selective Serotonin Reuptake Inhibitor Antidepressant Action: Systematic Review and Meta-analysis. Archives of General Psychiatry. 2006;63:1217–1223.

74. Brener J, Ring C. Towards a psychophysics of interoceptive processes: the measurement of heartbeat detection. Philosophical Transactions of the Royal Society B: Biological Sciences. 2016;371.

## Supplementary References

1. Montoya AK, Hayes AF. Two-condition within-participant statistical mediation analysis: A path-analytic framework. Psychol Methods. 2017;22:6–27.

2. Fleming SM, Ryu J, Golfinos JG, Blackmon KE. Domain-specific impairment in metacognitive accuracy following anterior prefrontal lesions. Brain. 2014;137:2811–2822.

3. Fleming SM, Lau HC. How to measure metacognition. Front Hum Neurosci. 2014;8.

4. Green DM, Swets JA. Signal detection theory and psychophysics. Oxford, England: John Wiley; 1966.

